# A reproducible and generalizable software workflow for analysis of large-scale neuroimaging data collections using BIDS Apps

**DOI:** 10.1101/2023.08.16.552472

**Authors:** Chenying Zhao, Dorota Jarecka, Sydney Covitz, Yibei Chen, Simon B. Eickhoff, Damien A. Fair, Alexandre R. Franco, Yaroslav O. Halchenko, Timothy J. Hendrickson, Felix Hoffstaedter, Audrey Houghton, Gregory Kiar, Austin Macdonald, Kahini Mehta, Michael P. Milham, Taylor Salo, Michael Hanke, Satrajit S. Ghosh, Matthew Cieslak, Theodore D. Satterthwaite

**Affiliations:** Lifespan Informatics and Neuroimaging Center (PennLINC), Department of Psychiatry, Perelman School of Medicine, University of Pennsylvania, Philadelphia, PA, USA; Penn/CHOP Lifespan Brain Institute, Perelman School of Medicine, Children’s Hospital of Philadelphia Research Institute, Philadelphia, PA, USA; Department of Bioengineering, School of Engineering and Applied Science, University of Pennsylvania, Philadelphia, PA, USA; Department of Psychiatry, Perelman School of Medicine, University of Pennsylvania, Philadelphia, PA, USA; McGovern Institute for Brain Research, Massachusetts Institute of Technology, Cambridge, MA, USA; Institute of Neuroscience and Medicine, Brain & Behaviour (INM-7), Research Center Jülich, Jülich, Germany; Institute of Systems Neuroscience, Medical Faculty, Heinrich Heine University Düsseldorf, Düsseldorf, Germany; Masonic Institute for the Developing Brain, University of Minnesota, Minneapolis, MN, USA; Institute of Child Development, College of Education and Human Development, University of Minnesota, Minneapolis, MN, USA; Department of Pediatrics, University of Minnesota Medical School, University of Minnesota, Minneapolis, MN, USA; Minnesota Supercomputing Institute, University of Minnesota, Minneapolis, MN, USA; Child Mind Institute, New York, NY, USA; Center for Biomedical Imaging and Neuromodulation, Nathan Kline Institute for Psychiatric Research, Orangeburg, NY, USA; Department of Psychiatry, NYU Grossman School of Medicine, New York, NY, USA; Department of Psychological and Brain Sciences, Dartmouth College, Hanover, NH, USA; Department of Otolaryngology, Harvard Medical School, Boston, MA, USA; Center for Biomedical Image Computation and Analytics, University of Pennsylvania, Philadelphia, PA, USA

**Keywords:** Reproducibility, BIDS Apps, software, MRI, big data, image processing

## Abstract

Neuroimaging research faces a crisis of reproducibility. With massive sample sizes and greater data complexity, this problem becomes more acute. Software that operates on imaging data defined using the Brain Imaging Data Structure (BIDS) – BIDS Apps – have provided a substantial advance. However, even using BIDS Apps, a full audit trail of data processing is a necessary prerequisite for fully reproducible research. Obtaining a faithful record of the audit trail is challenging – especially for large datasets. Recently, the FAIRly big framework was introduced as a way to facilitate reproducible processing of large-scale data by leveraging DataLad – a version control system for data management. However, the current implementation of this framework was more of a proof of concept, and could not be immediately reused by other investigators for different use cases. Here we introduce the <u>B</u>IDS <u>A</u>pp <u>B</u>oot<u>s</u>trap (BABS), a user-friendly and generalizable Python package for reproducible image processing at scale. BABS facilitates the reproducible application of BIDS Apps to large-scale datasets. Leveraging DataLad and the FAIRly big framework, BABS tracks the full audit trail of data processing in a scalable way by automatically preparing all scripts necessary for data processing and version tracking on high performance computing (HPC) systems. Currently, BABS supports jobs submissions and audits on Sun Grid Engine (SGE) and Slurm HPCs with a parsimonious set of programs. To demonstrate its scalability, we applied BABS to data from the Healthy Brain Network (HBN; n=2,565). Taken together, BABS allows reproducible and scalable image processing and is broadly extensible via an open-source development model.

## 1. INTRODUCTION

Lack of reproducibility in neuroscience – and neuroimaging in particular – has frequently been categorized as a “crisis”. While there are many facets to this crisis and barriers to reproducibility, one major source is analytic flexibility during image processing. Abundant analytic tools provide much greater flexibility in image processing. However, varying methods applied to the same dataset may lead to divergent results and conclusions (Botvinik-Nezer et al., 2020; Maier-Hein et al., 2017; Poldrack et al., 2017). These problems grow more acute as the size and complexity of imaging datasets increase. Large imaging data resources enhance statistical power (Button et al., 2013), and are often more diverse and representative (Laird, 2021), resulting in more generalizable findings. While there has been a well-motivated push to create large imaging data resources, most academic investigators who are end-users of these data resources lack essential tools for conducting reproducible research with large-scale imaging datasets. Here we introduce a user-friendly tool that facilitates fully reproducible processing of large-scale neuroimaging datasets.

Recent progress in reproducibility has been greatly facilitated by use of standardized data structures. The Brain Imaging Data Structure (BIDS) is the standard format for organizing brain imaging data from diverse modalities (structural, diffusion, functional images, etc) (Gorgolewski et al., 2016). BIDS includes not only the images, but also images’ metadata in sidecar JSON files (e.g., imaging parameters). BIDS Apps read such metadata to automatically configure the correct processing workflow, making them robust to heterogeneous data from different subjects and sessions. Although image processing software often depends on other software or packages, BIDS Apps are containerized (i.e., as Docker or Singularity container images) to encapsulate all dependencies and achieve portability even on different platforms, e.g., different high performance computing (HPC) clusters (Gorgolewski et al., 2017). Example BIDS Apps are QSIPrep (https://github.com/PennLINC/qsiprep; Cieslak et al., 2021) and fMRIPrep (https://github.com/nipreps/fmriprep; Esteban et al., 2019, 2020); similar BIDS Apps such as XCP-D (https://github.com/PennLINC/xcp_d; Adebimpe et al., 2023; Ciric et al., 2018) consume preprocessed output from BIDS Apps – or “BIDS derivatives” – and generate additional derived measures.

While containerized BIDS Apps provide a major advance for reproducible neuroscience, they do not automatically preserve a full audit trail along the way. Complete provenance of data processing results should answer *what* input data were, *which* version of the BIDS App was used, and *how* it was used (exact commands and parameters for running BIDS Apps, etc). Such a full audit trail of data processing is a necessary prerequisite for fully reproducible research. However, obtaining a faithful record of this audit trail is challenging – especially for large datasets that are processed using HPC clusters.

Version control tools provide a well-described way to record a full audit trail and enhance reproducibility. Git (https://git-scm.com/) has been used for code version control, however it is not efficient in tracking large binary data. Leveraging Git and git-annex (https://git-annex.branchable.com/), DataLad (https://www.datalad.org/; Halchenko et al., 2021) provides version control for data, even large binary files like neuroimaging data (e.g., in NIfTI format). The use of DataLad for large-scale reproducible image processing was introduced in the FAIRly big framework (Wagner et al., 2022). FAIR refers to findability, accessibility, interoperability, and reusability (Wilkinson et al., 2016). FAIRly big is a DataLad-based framework for reproducible processing of large-scale datasets (Wagner et al., 2022). DataLad and the FAIRly big framework capture provenance records of data processing, e.g., *who* applied *which* commands and code upon *which* input data to generate *which* output data. Such detailed records can be used for re-execution of the data processing and also provide a full audit trail (Wagner et al., 2022).

Although the FAIRly big framework paves the way for reproducible analysis at scale, its current implementation remains challenging. It requires investigators to write scripts for running the entire procedure; these scripts involve numerous steps and substantial proficiency with DataLad. This can be quite challenging for beginners – and is often hard to debug even for experienced users. In addition, these scripts also need to be customized for specific use cases. For example, the input datasets can include different data modalities (e.g., structural MRI, functional MRI, diffusion MRI); can be cross-sectional (single-session) studies or longitudinal (multiple-session) studies; and can be raw BIDS data or BIDS derivatives (e.g., fMRIPrep results). Furthermore, differences in the cluster systems also need to be accounted for. Different HPC job scheduling systems such as Sun Grid Engine (SGE) and Slurm often have different commands even for the similar functionality such as job submissions and status checking. One straightforward way to implement the FAIRly big framework is to write one script per use case, where input dataset, BIDS App, and cluster system are fixed. However, this approach results in a profusion of “one-off” programs due to the many combinations of various input dataset types, BIDS Apps and different cluster systems.

To address these challenges, we introduce <u>B</u>IDS <u>A</u>pp <u>B</u>oot<u>s</u>trap (BABS; http://pennlinc-babs.readthedocs.io/), a user-friendly and generalizable Python package for reproducible image processing at scale. Capitalizing on standard formats of neuroimaging data and containerized software, BABS takes BIDS datasets as input and applies BIDS Apps. The robustness of the BIDS App framework to data heterogeneity also facilitates the generalization of BABS to complex and large datasets. BABS automatically generates all code for data processing based on users’ customization, and records the full audit trail in a scalable way by leveraging DataLad and the FAIRly big framework. BABS’s automation and generalizability to different use cases are similar to those seen in BIDS Apps such as fMRIPrep and QSIPrep, which wrap up the entire preprocessing workflow and are generalizable to large, diverse neuroimaging datasets. With a parsimonious set of programs, BABS supports user-friendly job submissions and auditing on SGE and Slurm HPC clusters. As described below, BABS facilitates reproducible, generalizable, and scalable processing of BIDS datasets.

## 2. MATERIALS AND METHODS

### 2.1. Overview

BABS is a Python package for the reproducible application of BIDS Apps. It leverages DataLad and the FAIRly big framework to provide a full audit trail for the processing of large-scale datasets. It automatically “bootstraps” the need to execute the FAIRly big workflow: BABS generates all code for data processing and version tracking with DataLad. As part of this process, BABS interacts with HPC systems (e.g., SGE, Slurm) for submitting and auditing jobs. The entire BABS workflow can be completed with a parsimonious set of command-line programs.

### 2.2. Data provenance tracking via DataLad

BABS leverages DataLad (Halchenko et al., 2021) for data provenance tracking. Much like Git (https://git-scm.com/) provides version control for code, DataLad provides version control for data by building upon the functionality of Git and git-annex (https://git-annex.branchable.com/). Instead of directly tracking the file contents, DataLad tracks the “checksum” of the file content, a short, fixed-length hexadecimal number (e.g., 32 digits for MD5 checksum) representing the file content. This checksum can be used to verify the file content and check any changes in the file, as even a single byte change of the file’s content would result in a change of the checksum. Tracking this checksum is much cheaper than tracking the file content directly. Therefore, DataLad is capable of handling large files commonly seen in neuroimaging datasets. A DataLad dataset is a Git repository with a unique Universally Unique Identifier (UUID) and is destined for managing and tracking data if git-annex is enabled for that repository. As every change to the dataset can be recorded as a separate commit with comprehensive metadata, the ‘`datalad run`’ command can be used to save changes to the dataset as a result of the execution of any command, while recording the command within the Git commit associated with that change. Based on these, DataLad provides machine-readable, re-executable provenance records. The version-controlled DataLad dataset can also be cloned to another place for reuse or distribution. We refer the reader to the DataLad Handbook (http://handbook.datalad.org/; Wagner et al., 2023) to discover more about DataLad functionality and the larger ecosystem of extensions.

### 2.3. BABS workflow

BABS builds upon the workflow of the FAIRly big framework (see **Figure 1**), and can be used on HPC clusters. Here, HPC clusters are computing resources that include multiple connected servers, or “compute nodes”. HPC clusters utilize job scheduling systems to manage jobs running on different compute nodes. Thus, many jobs can be run in parallel on different compute nodes, making HPC clusters powerful tools for large-scale data processing. BABS supports SGE and Slurm, which are two of the most popular HPC job scheduling systems. BABS allocates the image processing for each subject (or session) into a “job” on HPC clusters. In this paper, we will follow the common principles in BIDS format and use the term “subject” to refer to a participant. The scope of a job is at subject-level for single-session dataset and at session-level for multiple-session dataset. For simplicity, in this paper, we may use “a specific session” to refer to “a specific session within a subject”. For each job, the input data from the corresponding subject (or session) will be cloned from the input Remote Indexed Archive (RIA) to an ephemeral (temporary) compute workspace. A RIA store is a permanent store, e.g., some storage space on HPC clusters. The input data is then processed by a BIDS App on a cluster compute node. The parameters for executing the BIDS App are predefined by the user in a YAML file; this will be explained in the next section “2.4. BABS programs”, as well as in the Results section with example YAML files. During the job run, all provenance is tracked by DataLad, including code (e.g., exact Singularity run command), input BIDS dataset(s), the BIDS App and its version, and the results. At the end of the job run, zipped results and the provenance are pushed to the output RIA store as a new branch. Processing for all subjects (or sessions) is parallelized. After all jobs are completed, results and provenance of successful jobs are merged and readily available in the output RIA store. Note that all the code for data processing and provenance tracking is automatically generated and internally executed – i.e., “bootstrapped” – by BABS.

**Figure 1.**
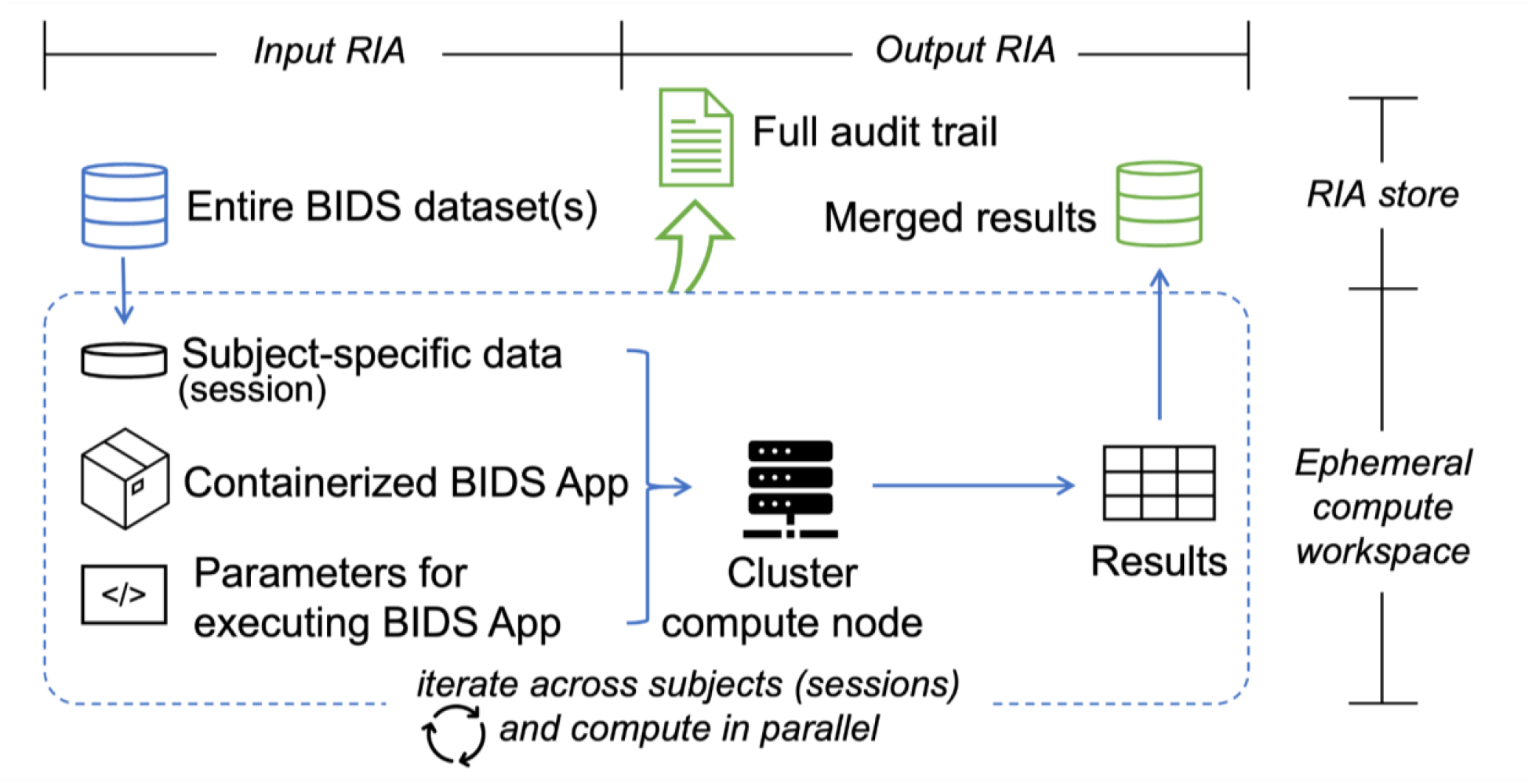
Schematic of the BABS workflow. For a job of a subject (single-session dataset) or a session (multiple-session dataset), the subject-(or session-) specific data is cloned from the permanent store – the input Remote Indexed Archive (RIA) store – to an ephemeral compute workspace, along with the containerized BIDS App and code for executing the BIDS App (left of the figure). The job will be completed at a cluster compute node. Jobs of all subjects (or sessions) are iteratively submitted to the compute nodes and computed in parallel (the entire box). Results from each job are zipped and pushed to the output RIA store as a separate branch (right of the figure). After all jobs have finished, results from all successful jobs are merged. The full audit trail of the successful jobs is also saved in the output RIA store (top of the figure).

### 2.4. BABS programs

To achieve steps in the BABS workflow, BABS features several command-line interface programs (**Figure 2**). These programs can be run in a terminal connected to an HPC cluster. Detailed descriptions of how to use these programs can be found in the online documentation: https://pennlinc-babs.readthedocs.io/en/stable/cli.html.

**Figure 2.**
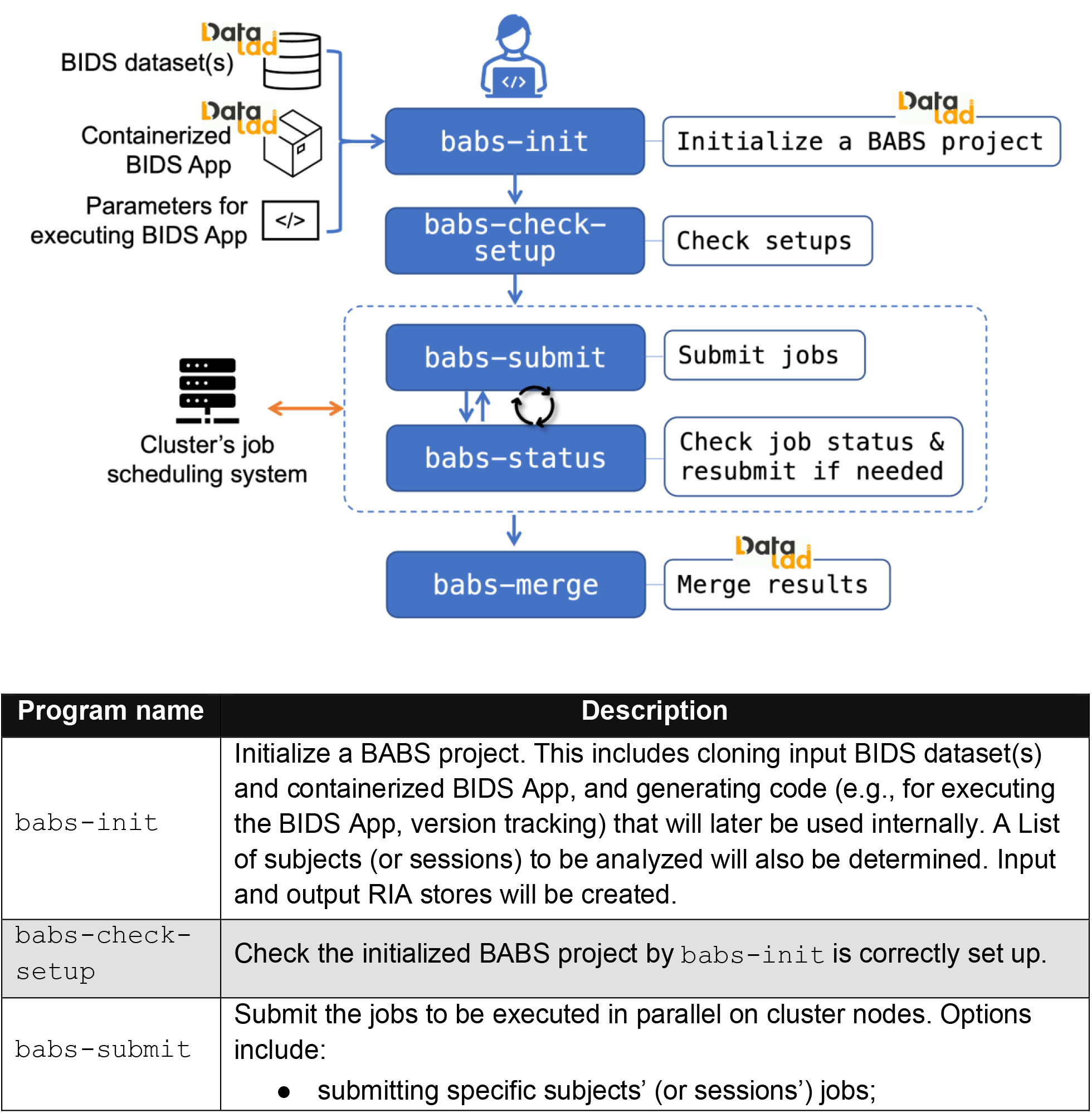
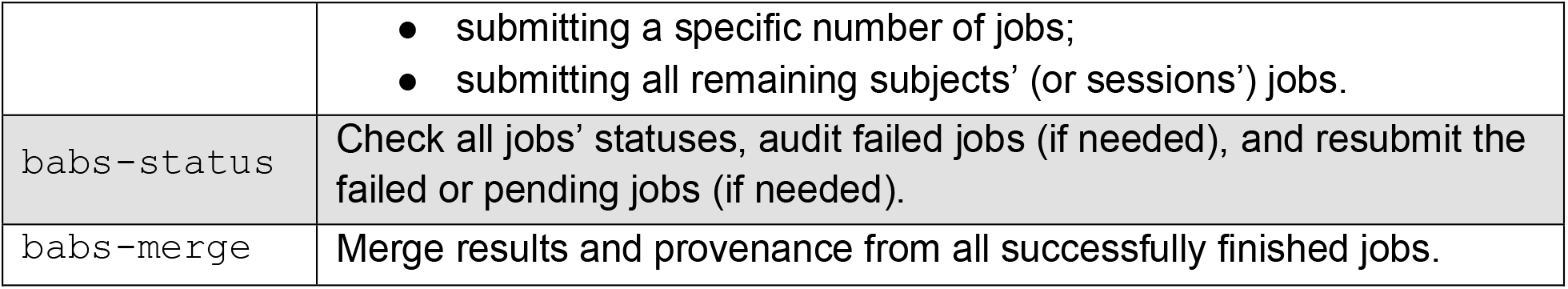
BABS user-oriented workflow (top figure) and descriptions of BABS programs (bottom table).

BABS requires three inputs: (a) one or more BIDS DataLad datasets, (b) a DataLad dataset of containerized BIDS App, and (c) a container configuration YAML file. Instead of directly using BIDS datasets or BIDS App images, we require them to be tracked by DataLad, i.e., as “DataLad datasets”, so that these two inputs to BABS come with their version history. DataLad datasets of BIDS and containerized BIDS App can also be easily cloned by BABS. Here, a “container DataLad dataset” means a collection of container images in a folder tracked by DataLad. As the content is a set of container images, it contains software that is used in the image processing workflow. It should be noted that BABS also supports input BIDS DataLad datasets that are remote, e.g., on OSF (e.g., see “3.2. Example walkthrough” in the Results section), or on some local servers to which HPC compute nodes have access. If the HPC compute nodes have access to the remote file system or the internet, users can simply provide the path to the remote BIDS DataLad dataset as the input to BABS. In this case, users do not need to download all the data content into the permanent disk space of HPC clusters before running BABS. Instead, when jobs are running on compute nodes, the scripts generated by BABS will utilize DataLad programs and automatically download the data content needed for each job into the ephemeral (temporary) compute workspace.

The last required input, a container configuration YAML file, is used to define how the BIDS App should be executed. Example YAML files are shown in **Figure 3A** and **Figure 4**. This YAML file is designed to be abstracted from the specifics of different cluster system types (e.g., SGE, Slurm). For example, there are several commonly used directive commands for requesting cluster resources in SGE and Slurm. Although they have similar goals (e.g., requesting memory), the commands are different in SGE and Slurm clusters. To reduce the differences in YAML files for different clusters, several commonly used directive commands that share similar functions on SGE and Slurm have been abstracted into keywords, e.g., `hard_memory_limit` (see below: line #14 in **Figure 3A**; line #22 in **Figure 4**). Most of these keywords can be used for both SGE and Slurm clusters without further changes. This facilitates the reuse of YAML files on another cluster with only minor changes. It should be noted that it is unavoidable that differences between specific HPC clusters will require some minor customizations of the YAML files, e.g., how the execution environment is configured. We provide some examples on how to customize the YAML files in the “3.2. Example walkthrough” in the Results section. More details can be found in BABS documentation: https://pennlinc-babs.readthedocs.io/en/stable/preparation_config_yaml_file.html.

**Figure 3.**
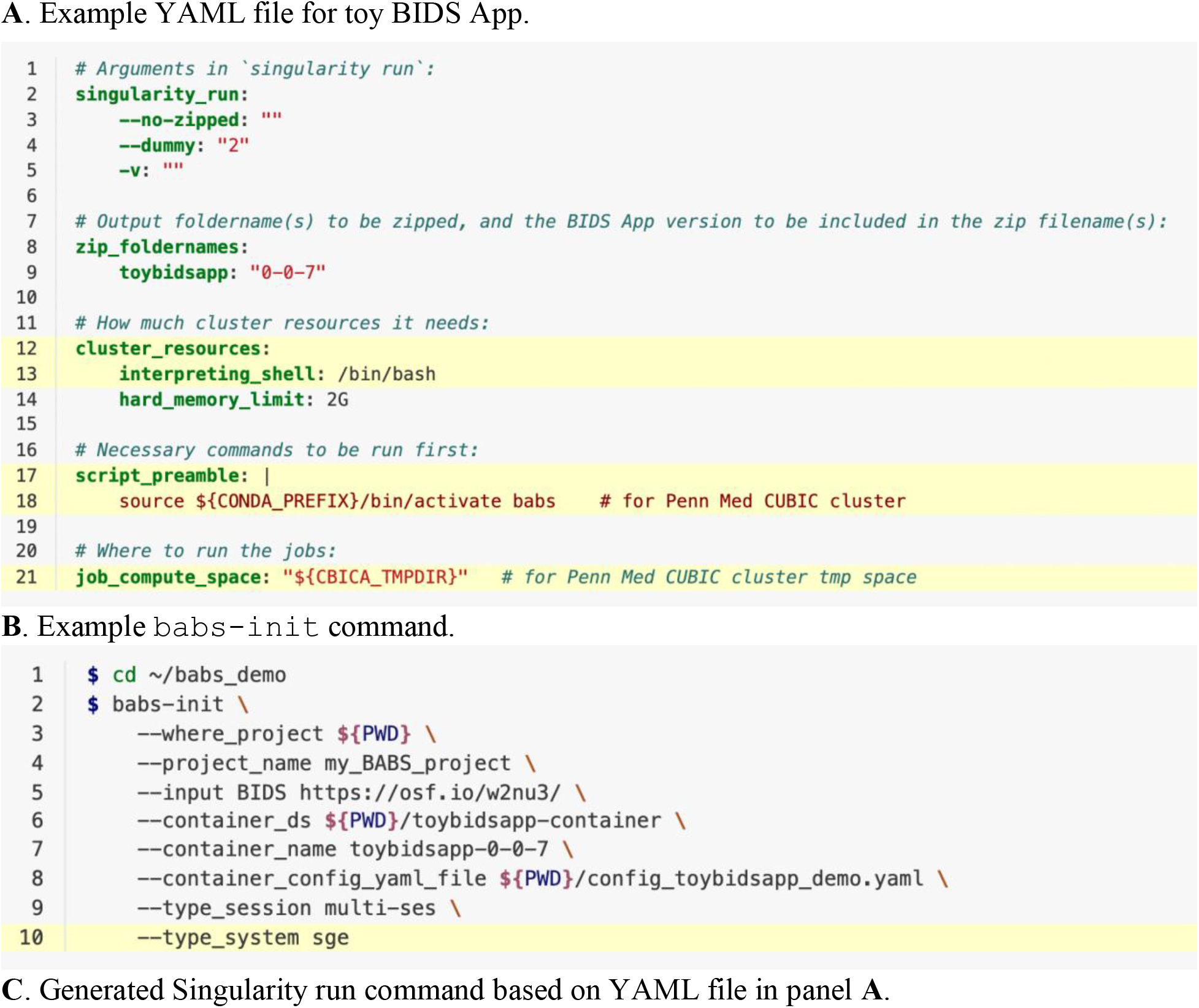
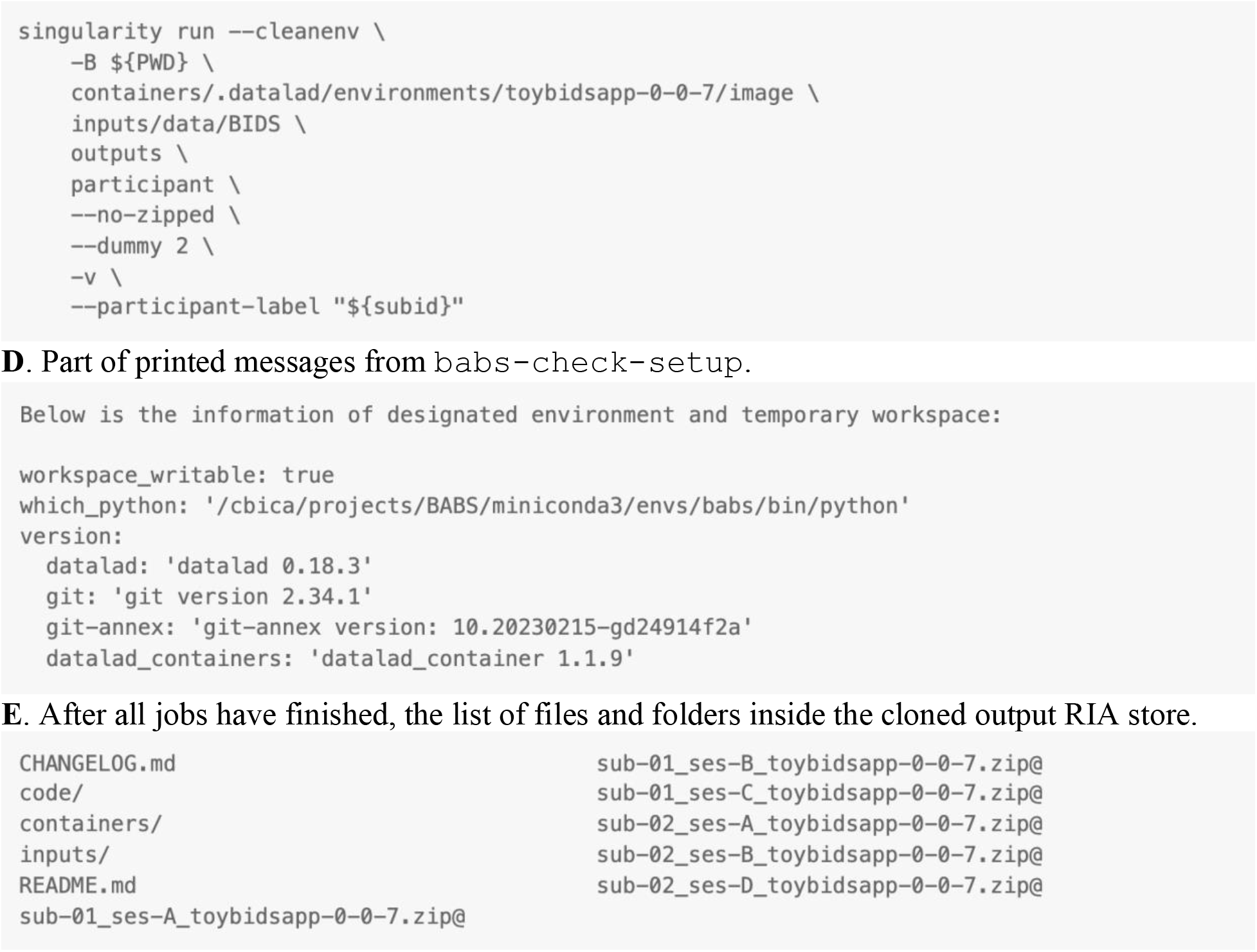
Materials used and generated in the example walkthrough of BABS. (**A**) Example YAML file for toy BIDS App. This YAML file was prepared for Penn Medicine CUBIC SGE cluster, however, with some customization (highlighted lines) based on the clusters users are using, this YAML file can also be applied to other clusters, even for a Slurm cluster. (**B**) Example `babs-init` command. The highlighted line requires customization. (**C**) Generated Singularity run command based on the YAML file in panel **A**. (**D**) Part of printed messages from `babs-check-setup`, which provides information of designated environment and temporary job compute space. (**E**) After all jobs have finished, the list of files and folders inside the cloned output RIA store.

**Figure 4.**
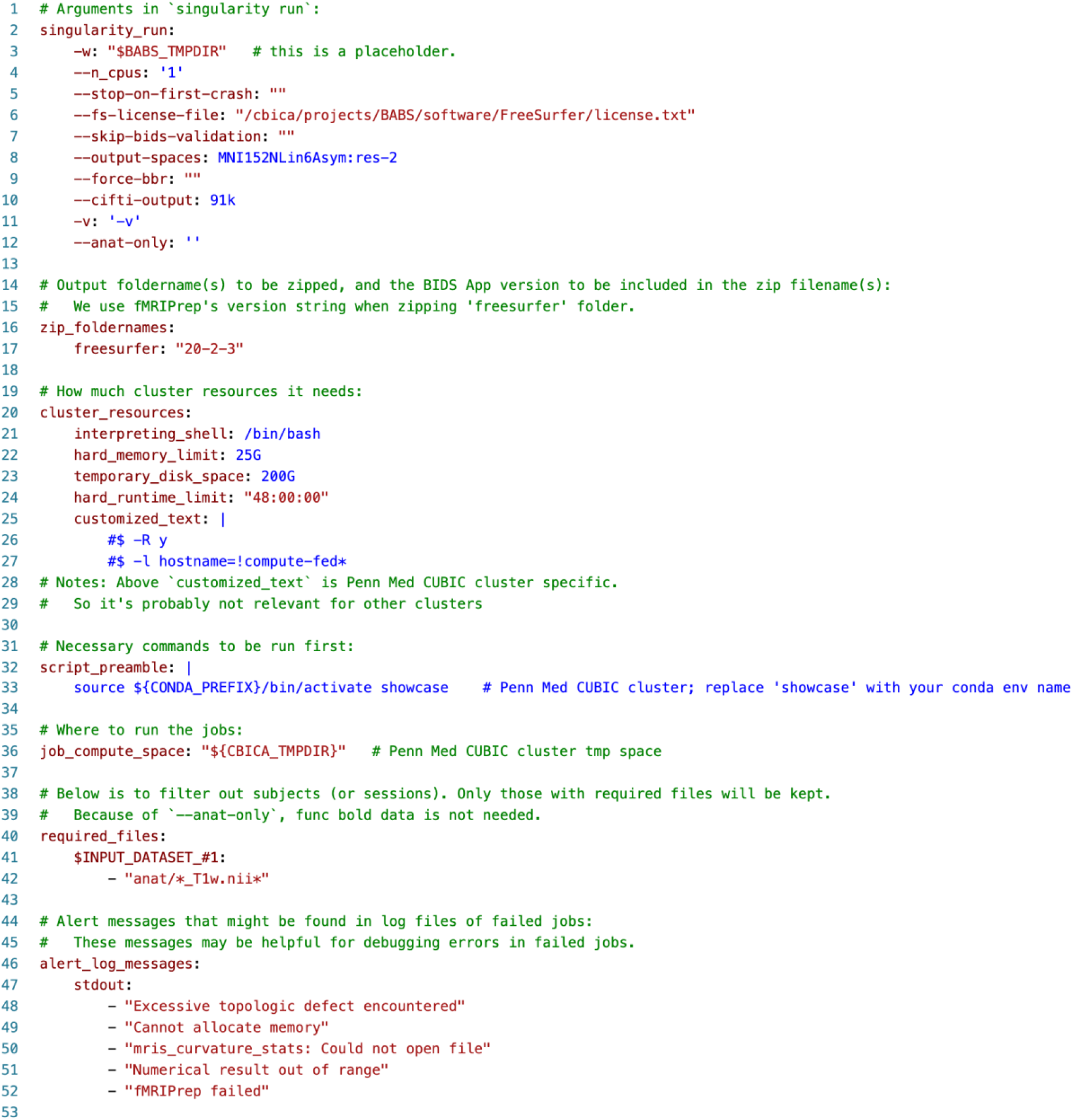
Container configuration YAML file used in application of BABS to the HBN dataset.

With these three required inputs, `babs-init` initializes a BABS project, a folder that will hold input data, container, code, and results. A BABS project can be located in the storage space of a cluster. `babs-init` will create an analysis folder within this BABS project, which will hold input BIDS dataset(s), container DataLad dataset, code, etc. Importantly, this analysis folder is itself a DataLad dataset whose nested content – including data and code – is tracked by DataLad. `babs-init` then creates input and output RIA stores as the DataLad siblings (“copies”) of analysis DataLad dataset. `babs-init` clones the input BIDS DataLad dataset(s) and the container DataLad dataset into the analysis DataLad dataset. In addition, it generates the code to be used internally, including code for executing the desired BIDS App and DataLad version control. `babs-init` also determines the list of the subjects (or sessions) to analyze based on an initial inclusion list optionally provided by the user (argument `--list-sub-file`). This list can be further filtered by excluding those subjects (or sessions) that do not have required files optionally defined in the container configuration YAML file.

After initializing a BABS project, `babs-check-setup` will be used to confirm this BABS project has been set up properly. Specifically, it will perform sanity checks of the components of this BABS project, e.g., if all necessary scripts have been generated, if input datasets have been successfully cloned, etc. We highly recommend users submit a toy job using argument `--job-test` to make sure necessary packages (e.g., DataLad) are installed in the designated environment, and that setup specified in the container configuration YAML file (e.g., section `script_preamble`) is working as expected. `babs-check-setup` can be also used as a diagnostic tool, as it prints out information for users to review, including configurations of the BABS project (e.g., input BIDS dataset’s name and path) and versions of necessary packages in the designated environment. This information is also helpful for submitting bug reports if issues arise.

Once the setup is complete, the BABS project is ready for job submitting and status monitoring via `babs-submit` and `babs-status`. These two programs interact with the cluster’s job scheduling system and can be used iteratively. Each job of data processing operates on a specific subject or a specific session. `babs-submit` provides several submission options, including submitting specific subjects’ (or sessions’) jobs, submitting a specific number of jobs, and submitting all the remaining jobs.

`babs-status` can be used to check job status. The job status of a subject or a session can be one of these categories: (a) the job has not been submitted yet; (b) the job has been submitted but is waiting in the queue (“pending”); (c) the job has been submitted and is running on a compute node; (d) the job has successfully finished; (e) the job failed with an error. `babs-status` will check job statuses from all subjects (or sessions) and print out a summary containing the number of jobs in each category. In addition, `babs-status` can also perform auditing on failed jobs to provide more information regarding why those jobs were failed. Finally, `babs-status` can also be used to resubmit failed or pending jobs. Options include resubmitting specific subjects’ (or sessions’) jobs, and resubmitting all of failed or pending jobs. Note that although the processing of a subject’s (or session’s) images can be resubmitted using `babs-status`, at any given time, there should be only one job submitted and under “running” or “pending” for a specific subject (or session).

Results from each job are compressed (“zipped”) and are kept in a separate branch. After all jobs are finished, `babs-merge` can be used to merge all the results and provenance from successfully finished jobs into the mainline branch in the output RIA. After merging, results are ready to consume. Users can use DataLad commands to clone the output RIA, get the results content, and unzip the results.

### 2.5. Open-source software development and release

The source code of BABS is version controlled and publicly available on GitHub (https://github.com/PennLINC/babs). We have been using CircleCI to run tests and make releases. Specifically, each new commit pushed to GitHub triggers CircleCI jobs to run the BABS’s unit tests. This helps ensure the quality and stability of BABS after each commit. When a version tag is pushed to GitHub, a new version of the Python package of BABS will be automatically built by CircleCI jobs and publicly released on the Python Package Index (PyPI): https://pypi.org/project/babs/.

### 2.6. Data and code availability statement

BABS documentation can be found at: http://pennlinc-babs.readthedocs.io/. Code for preparing application of BABS to the Healthy Brain Network (HBN) dataset is available at https://github.com/PennLINC/babs_paper. The source code of BABS is available at https://github.com/PennLINC/babs. The Python package of BABS can be downloaded from PyPI: https://pypi.org/project/babs/. The version of BABS used for preparing the example walkthrough and used in the application to the HBN dataset was 0.0.3. The container image of the toy BIDS App used in the example walkthrough is available on Docker Hub: https://hub.docker.com/r/pennlinc/toy_bids_app. The toy BIDS dataset used in the example walkthrough is available on OSF: https://osf.io/w2nu3/. Part of the HBN dataset is available on FCP-INDI (https://fcp-indi.s3.amazonaws.com/index.html#data/Projects/HBN/BIDS_curated/) (Richie-Halford et al., 2022).

### 2.7. Ethics statement

No new data were collected specifically for this paper. The Healthy Brain Network (HBN; Alexander et al., 2017) study was approved by the Chesapeake Institutional Review Board. Informed consent was obtained from each participant aged 18 or older. For participants younger than 18, written consent was obtained from their legal guardians and written assent was obtained from the participant. In addition, during the consent process, all participants provide informed consent for their data to be shared via IRB approved protocols.

## 3. RESULTS

### 3.1. Example use cases

BABS is designed to process large BIDS datasets with BIDS Apps on HPC clusters with job scheduling systems. Currently, BABS supports two popular HPC job scheduling systems, SGE and Slurm. The major differences between job scheduling systems lie in different commands for how the jobs are managed, e.g., in job submissions, status checking, etc. BABS generates and uses different code and commands of job managements tailored for different job scheduling systems.

BABS can be used to process data with BIDS Apps. Thus far, BABS has been used to process data using fMRIPrep (https://github.com/nipreps/fmriprep; Esteban et al., 2019, 2020), QSIPrep (https://github.com/PennLINC/qsiprep; Cieslak et al., 2021), and XCP-D (https://github.com/PennLINC/xcp_d; Ciric et al., 2018) (**Table 1**). Besides the BIDS Apps listed in **Table 1**, we also provide a toy BIDS App for quickly testing BABS (https://hub.docker.com/r/pennlinc/toy_bids_app). This toy BIDS App will be used in the example walkthrough (see below).

**Table 1.** Example use cases of BABS. fMRI, functional MRI; dMRI, diffusion MRI.

| # | Use cases / BIDS Apps | 1st input BIDS dataset | 2nd input BIDS dataset |
| --- | --- | --- | --- |
| 1 | fMRIPrep (for fMRI) | Raw BIDS (unzipped) | N/A |
| 2 | fMRIPrep (with FreeSurfer results ingressed) | Raw BIDS (unzipped) | FreeSurfer results (BIDS derivatives, zipped) |
| 3 | QSIprep (for dMRI) | Raw BIDS (unzipped) | N/A |
| 4 | XCP-D (for fMRI) | fMRIPrep results (BIDS derivatives, zipped) | N/A |

BABS accepts different input BIDS datasets, including raw BIDS datasets, or BIDS derivatives datasets. The latter case often includes results from another BIDS App, e.g., FreeSurfer results from fMRIPrep anatomical workflow. When using such BIDS derivatives as input datasets, currently BABS expects that the results of each subject (or session) are zipped. Therefore, we also refer to it as a “zipped BIDS derivatives dataset”. BABS also allows more than one input BIDS dataset. An example use case would be applying the fMRIPrep BOLD preprocessing workflow upon functional MRI data from a raw BIDS dataset, with another input BIDS derivatives dataset, FreeSurfer results, ingressed (**Table 1**, the second use case). For use cases listed in **Table 1**, we have provided example container configuration files available in BABS GitHub repository. Please refer to this markdown file for the list of available files and their links: https://github.com/PennLINC/babs/blob/main/notebooks/README.md.

Beyond the existing tested applications in **Table 1**, BABS can be used to process data with most BIDS Apps after users create new container configuration YAML files for these BIDS Apps accordingly. Furthermore, BABS can be extended to be used with other scheduling systems (e.g., LSF) that are not yet supported. We welcome enhancements from the user community via new pull requests at the BABS GitHub repository (https://github.com/PennLINC/babs). Below, we demonstrate the use of BABS first via a detailed walkthrough using a toy example. Second, we illustrate its application to a large-scale dataset (the structural data of the Healthy Brain Network).

### 3.2. Example walkthrough

To demonstrate an example usage of BABS, we provide an example walkthrough where a toy BIDS dataset and toy BIDS App are used. The example walkthrough is also available as online documentation: https://pennlinc-babs.readthedocs.io/en/stable/walkthrough.html, where more details are provided. We recommend referring to the online version of this example walkthrough and copying and pasting the commands from there to ensure use of the most current version and to allow for tighter control over command text formatting (compared to a journal publication). As detailed below, there are four major steps in this example walkthrough: (1) Get prepared; (2) Create a BABS project; (3) Submit jobs and check job status; (4) After jobs have finished.

### Step 1. Get prepared

We first install BABS and dependent packages in a conda environment called `babs`. The installation steps have been detailed in our online documentation: https://pennlinc-babs.readthedocs.io/en/stable/installation.html. Note that besides the required dependencies like DataLad, Git, git-annex, and datalad-container, we also installed datalad-osf Python package so that we can use a toy BIDS dataset available on OSF as input. We used BABS version 0.0.3 to prepare this example walkthrough. As there would be more enhancements in BABS in future releases that might potentially alter the commands or steps here, we encourage users to use the latest BABS version available on PyPI and the latest stable version of walkthrough available online (https://pennlinc-babs.readthedocs.io/en/stable/walkthrough.html).

We first create a folder called babs_demo in the root directory as the working directory in this example walkthrough:

~~~
conda activate babs
mkdir -p ∼/babs_demo
cd babs_demo
~~~

Now, we need to prepare three inputs required by BABS: an input BIDS DataLad dataset, a DataLad dataset of containerized BIDS App (“container DataLad dataset”), and a container configuration YAML file. For the input BIDS DataLad dataset, we can use a toy, multiple-session BIDS DataLad dataset publicly available on OSF: https://osf.io/w2nu3/. In this BIDS dataset, there are two subjects, each with three sessions. As long as the compute nodes of the HPC cluster are connected to the internet, the OSF link of this dataset can be directly copied as the path to the input BIDS DataLad dataset when running `babs-init`, and BABS will download all necessary files. For compute nodes without internet access, please refer to the online version of the walkthrough: https://pennlinc-babs.readthedocs.io/en/stable/walkthrough.html.

For the BIDS App, we will use the toy BIDS App in this example walkthrough to allow for very rapid execution of the jobs. For a raw BIDS dataset like the one we will use in this walkthrough, this toy BIDS App performs a simple task when used with BABS: it will count non-hidden files in a subject’s folder. Its Docker image is publicly available on Docker Hub: https://hub.docker.com/r/pennlinc/toy_bids_app. To prepare its container DataLad dataset, we first pull toy BIDS App (version 0.0.7) as a Singularity image from the Docker Hub:

~~~
cd ∼/babs_demo
singularity build \
  toybidsapp-0.0.7.sif \
  docker://pennlinc/toy_bids_app:0.0.7
~~~

Now we can see the Singularity image file `toybidsapp-0.0.7.sif` in the current directory. We then create a DataLad dataset of this container (i.e., let DataLad track this Singularity image):

~~~
datalad create -D “toy BIDS App” toybidsapp-container
cd toybidsapp-container
datalad containers-add \
  --url ${PWD}/../toybidsapp-0.0.7.sif \
  toybidsapp-0-0-7
~~~

Now, the DataLad dataset `toybidsapp-container` which contains the toy BIDS App container is ready to use. As the Singularity image file has been copied into `toybidsapp-container`, we can remove the original Singularity image file:

~~~
cd ..
rm toybidsapp-0.0.7.sif
~~~

Finally, we need to prepare a YAML file that instructs BABS for how to run the BIDS App. **Figure 3A** shows an example YAML file for toy BIDS App, and we will use it in this example walkthrough. Note that this YAML file was prepared for Penn Medicine CUBIC SGE cluster, however, this YAML file can also be applied to other clusters (including Slurm clusters) after some customization (highlighted lines).

There are several sections in this YAML file. The first section `singularity_run` defines the arguments and their values for running the BIDS App (line #2-5, **Figure 3A**). The argument `--no-zipped` tells the toy BIDS App that the input dataset is unzipped, raw BIDS dataset. The other two arguments, `--dummy` and `-v` are both examples of what an argument could look like, where argument `--dummy` can take any value afterwards, and argument `-v` does not take values.

The next section, `zip_foldernames`, is about the results zip files (line #8-9, **Figure 3A**). As the results from each subject (or each session) will be zipped, here we tell BABS the name(s) of the output folder(s) to be zipped is `toybidsapp`. In addition, we also provide the version of this toy BIDS App, `0-0-7`, so that it can be added to the zip filenames. This will result in the zip filenames being named according to the convention of `sub-01_ses-A_toybidsapp-0-0-7.zip` for subject 01 (`sub-01`) and session A (`ses-A`).

The third section, `cluster_resources` (line #12-14, **Figure 3A**), defines cluster resource requirements such as the memory requirement (line #14, **Figure 3A**) using scheduler-agnostic keywords. The interpreting shell to be used in the job script (line #13, **Figure 3A**) is also defined in this section. These items will be converted to scheduler-specific directives in the job script.

There are inevitable idiosyncrasies across clusters, thus, this section often requires customization by users for their clusters. For the line of `interpreting_shell`, some Slurm clusters might suggest users to use ‘`interpreting_shell: “/bin/bash −l”`’ instead, however, users should consult their clusters’ documentation and administrator. Customized commands can also be added after `customized_text`, without using the predefined cluster resources keywords in BABS. For example, for Slurm clusters, users may request specific partition(s) via:

~~~
cluster_resources:
 …
 customized_text: |
  #SBATCH -p
~~~

The fourth section, `script_preamble` (line #17-18, **Figure 3A**), defines commands that should be run before data processing starts. It could include commands for setting up the virtual environment (line #18, **Figure 3A**), for loading necessary modules, etc. Because each cluster may be configured differently, this section often requires customization.

The final section in this YAML file is called `job_compute_space` (line #21, **Figure 3A**). This is to set the location of the compute space where the jobs will run. As results will be saved to the permanent storage space output RIA, we recommend using a temporary space here, such as space on the compute node, to avoid risk of accumulating unnecessary data from failed jobs which takes up space. The path `“${CBICA_TMPDIR}”` in line #21 in **Figure 3A** was specifically used for Penn Medicine CUBIC cluster, and other clusters will likely have different paths to the temporary compute space, so customization is needed here.

Once the user has finished inputting all necessary customization, we save the YAML file as file `config_toybidsapp_demo.yaml` into directory `∼/babs_demo`. Note that currently this directory also includes the container DataLad dataset `toybidsapp-container`.

### Step 2. Create a BABS project

With all three inputs ready, we can now start to use BABS for data analysis. We first use `babs-init` to create a BABS project. This is a folder where input DataLad datasets of BIDS dataset(s) and the containerized BIDS App are cloned to, all scripts are generated, and results and provenance are saved. **Figure 3B** shows an example command of `babs-init`. With this example command, we create a BABS project called `my_BABS_project` (line #4, **Figure 3B**) in directory `∼/babs_demo`. We call the input dataset as `BIDS`, and we provide the OSF link as its path (line #5, **Figure 3B**). For the container and its execution, we use the container DataLad dataset `toybidsapp-container` and the YAML file we just prepared (line #6-8, **Figure 3B**). We make sure that the string `toybidsapp-0-0-7` used in `--container_name` (line #7, **Figure 3B**) is consistent with the image name we specified when preparing `toybidsapp-container`. As this input BIDS dataset is a multiple-session dataset, we specify this as `‘--type_session multi-ses’` (line #9, **Figure 3B**). Finally, because we will run this on an SGE cluster, we specify the cluster system type as `’--type_system sgè` (highlighted line #10, **Figure 3B**). If a Slurm cluster is used, a user would change line #10 to `‘--type_system slurm’`. After running this `babs-init` command, we see this message at the end, indicating the success: “`‘babs-init’ was successful!`”.

It is very important to check two things in the generated code. The first is the Singularity run command for running the BIDS App. This command has been printed out by `babs-init` (see **Figure 3C**). As you can see, the arguments and their values specified in the `singularity_run` section in YAML file (line #2-5, **Figure 3A**) has been added to the generated Singularity run command (**Figure 3C**). BABS has also automatically handled the positional arguments of the BIDS App, including input directory, output directory, and analysis level (`’participant’`). The `--participant-label` parameter is also covered by BABS.

The second thing to check is the generated directives in the job script. These directives are in the header lines of the script. We get them via:

~~~
cd ∼/babs_demo/my_BABS_project
head analysis/code/participant_job.sh
~~~

The first several lines starting with ‘`#`’ and before the line ‘`# Script preambles:`’ are the generated directives. Using the YAML file above without further modifications, for BABS version > 0.0.3 applied on an SGE cluster, we will see these directives:

~~~
#!/bin/bash
#$ -l h_vmem=2G
~~~

and on an Slurm cluster:

~~~
#!/bin/bash
#SBATCH --mem=2G
~~~

At this point, a folder named `my_BABS_project` has been generated in the directory `∼/babs_demo`. This folder includes three sub-folders, `analysis`, `input_ria`, and `output_ria`. The folder `analysis` is also a DataLad dataset which includes the cloned inputs (input BIDS DataLad dataset and container DataLad dataset), and generated scripts. The folders `input_ria` and `output_ria` are the input and output RIA stores, respectively, and they are DataLad siblings of `analysis`. When jobs are running, inputs are cloned from input RIA store, and results and provenance will be pushed to output RIA store.

It is important to let BABS check to be sure that the project has been initialized correctly before attempting to run many data processing jobs. One should run a test job to make sure that the environment and cluster resources specified in the YAML file are workable. We use `babs-check-setup` to do so. Note that the following BABS commands will be called from where the BABS project is located, `∼/babs_demo/my_BABS_project`. After switching to this directory, we can use `${PWD}` for argument `--project-root` in BABS commands.

~~~
cd ∼/babs_demo/my_BABS_project
babs-check-setup \
  --project-root ${PWD} \
  --job-test
~~~

Because we ask `babs-check-setup` to submit a test job, it might take a bit of time for the above command to finish, depending on how busy the cluster is. After running `babs-check-setup`, we see this message at the end, indicating the success: “`‘babs-check-setup’ was successful!`”.

Before moving on, we review the summarized information of the designated environment and temporary compute space where the jobs will run. This summarized information has been printed out by `babs-check-setup` (see **Figure 3D**). We confirm that the temporary compute space is writable (`‘true’`), the Python interpreter is what we desire, and the required packages have been installed, and their version numbers are appropriate.

### Step 3. Submit jobs and check job status

We will iteratively use `babs-submit` and `babs-status` to submit jobs and check job status. We first use `babs-status` to check the number of jobs we initially expect to finish successfully. In this example walkthrough, as no initial list was provided, BABS determines this number based on the number of sessions in the input BIDS dataset. We did not request extra filtering (based on required files) in our YAML file either, so BABS will submit one job for each session.

~~~
cd ∼/babs_demo/my_BABS_project
babs-status --project-root $PWD
~~~

Printed messages from babs-status tell us that “There are in total of 6 jobs to complete”.

We now use `babs-submit` to submit one job and see if it will finish successfully. By default, `babs-submit` will only submit one job.

~~~
babs-submit --project-root $PWD
~~~

Now, the job for the first session, `sub-01`/`ses-A` has been submitted. We can check the job status via `babs-status`:

~~~
babs-status --project-root $PWD
~~~

If this first job finished successfully, the printed messages from babs-status will tell us that “1 job(s) are successfully finished”.

Now, we submit all other jobs by specifying --all:

~~~
babs-submit --project-root $PWD --all
~~~

We can again call ‘`babs-status --project-root $PWD’` to check status. `babs-status` will tell us the number of jobs submitted, finished, pending, running, or failed. If all jobs have finished successfully, we will see printed messages: “`6 job(s) are successfully finished`” and “`All jobs are completed!`”.

### Step 4. After jobs have finished

After all jobs have finished successfully, we will merge all the results and provenance. Each job was executed on a different branch, so we must merge them together into the mainline branch. We now run `babs-merge` in the root directory of `my_BABS_project`:

~~~
babs-merge --project-root $PWD
~~~

After this command finishes running, we see “`’babs-mergè was successful!`” at the end of the printed messages. Now we are ready to consume the results.

To consume the results, we should not access the output RIA store or `merge_ds` directories inside the BABS project. Instead, we will clone the output RIA as another folder (e.g., called `my_BABS_project_outputs`) to a location external to the BABS project:

~~~
cd ..
datalad clone \
  ria+file://${PWD}/my_BABS_project/output_ria#∼data \
  my_BABS_project_outputs
~~~

The first command, ‘`cd ..`’ changes the directory back to folder `babs_demo`, where `my_BABS_project` locates. After running above commands, we then go into this new folder `my_BABS_project_outputs` and see what is inside:

~~~
cd my_BABS_project_outputs
ls
~~~

The content in this folder is shown in **Figure 3E**. As we see, results of each session have been saved in a zip file. Before unzipping a zip file, e.g., the zip file for `sub-01`/`ses-A`, we need to get its content first:

~~~
datalad get sub-01_ses-A_toybidsapp-0-0-7.zip
unzip sub-01_ses-A_toybidsapp-0-0-7.zip
~~~

From the zip file, we get a folder called `toybidsapp`. We check the content of this folder via:

~~~
cd toybidsapp
ls
~~~

In this folder, there is a file called `num_nonhidden_files.txt`. This is the result from the toy BIDS App, which is the number of non-hidden files in this subject’s data. Note that for raw BIDS dataset and when used with BABS, toy BIDS App counts at subject-level, even though current dataset is a multiple-session dataset. We print out the content of this text file:

~~~
cat num_nonhidden_files.txt
~~~

And the content should be “67”. “67” is the expected number for `sub-01` (which we are looking at), “56” is the expected number for `sub-02`. Getting the expected number means that toy BIDS App and BABS ran as expected.

### 3.3. Application to a large-scale dataset

Next, we applied BABS to the HBN dataset using the fMRIPrep anatomical workflow. As above, this was executed on the Penn Medicine CUBIC SGE cluster. HBN is a single-session dataset, i.e., each subject only has one session. We first prepared the input BIDS DataLad dataset, container DataLad dataset of fMRIPrep (Esteban et al., 2019; version 20.2.3), and container configuration YAML file (see **Figure 4**). In this container configuration YAML file, we set the job run time limit of two days (line #24, **Figure 4**), which is sufficient for fMRIPrep anatomical workflow. Jobs which could not finish within two days are probably stuck and should be killed by the cluster.

We first ran `babs-init` to initialize the BABS project. After it finished, we used `babs-check-setup` with `--job-test` mode on to make sure that the environment and cluster resources specified in the YAML file were workable. `babs-check-setup` command finished successfully, suggesting that real jobs of fMRIPrep were ready to submit.

Next, we used `babs-status` to check how many subjects’ jobs there are to complete. The list of subjects was generated during `babs-init`, where only subjects with T1-weighted images were included, and such data filtering was set on line #40-42 in the YAML file shown in **Figure 4**. There were 2,565 jobs to complete, where each job corresponds to a subject. We then submitted the first ten jobs using `babs-submit` with argument ‘`--count 10`’ to confirm real jobs could finish without error. After almost a day, all of these ten jobs finished successfully according to `babs-status`. This told us that the requested cluster resources were sufficient for the real jobs.

We then submitted jobs for all of the remaining subjects using `babs-submit` with argument `--all`. After that, we ran `babs-status` episodically to check job status. **Figure 5** shows example printed messages from `babs-status` during this process. The `babs-status` command is highlighted in blue. To audit the failed jobs, we used container configuration YAML file (line #3-4, **Figure 5**) and turned on `--job-account` mode (line #5, **Figure 5**). There were two major parts in the printed messages: the job status summary and the failed job audits. Based on the job status summary, at that point, all 2,565 subjects’ jobs had been submitted (line #11, **Figure 5**). Among them, 1,808 jobs successfully finished; 196 jobs were waiting in the queue (“pending”); 342 jobs were running; and 219 jobs failed (line #12-16, **Figure 5**).

**Figure 5.**
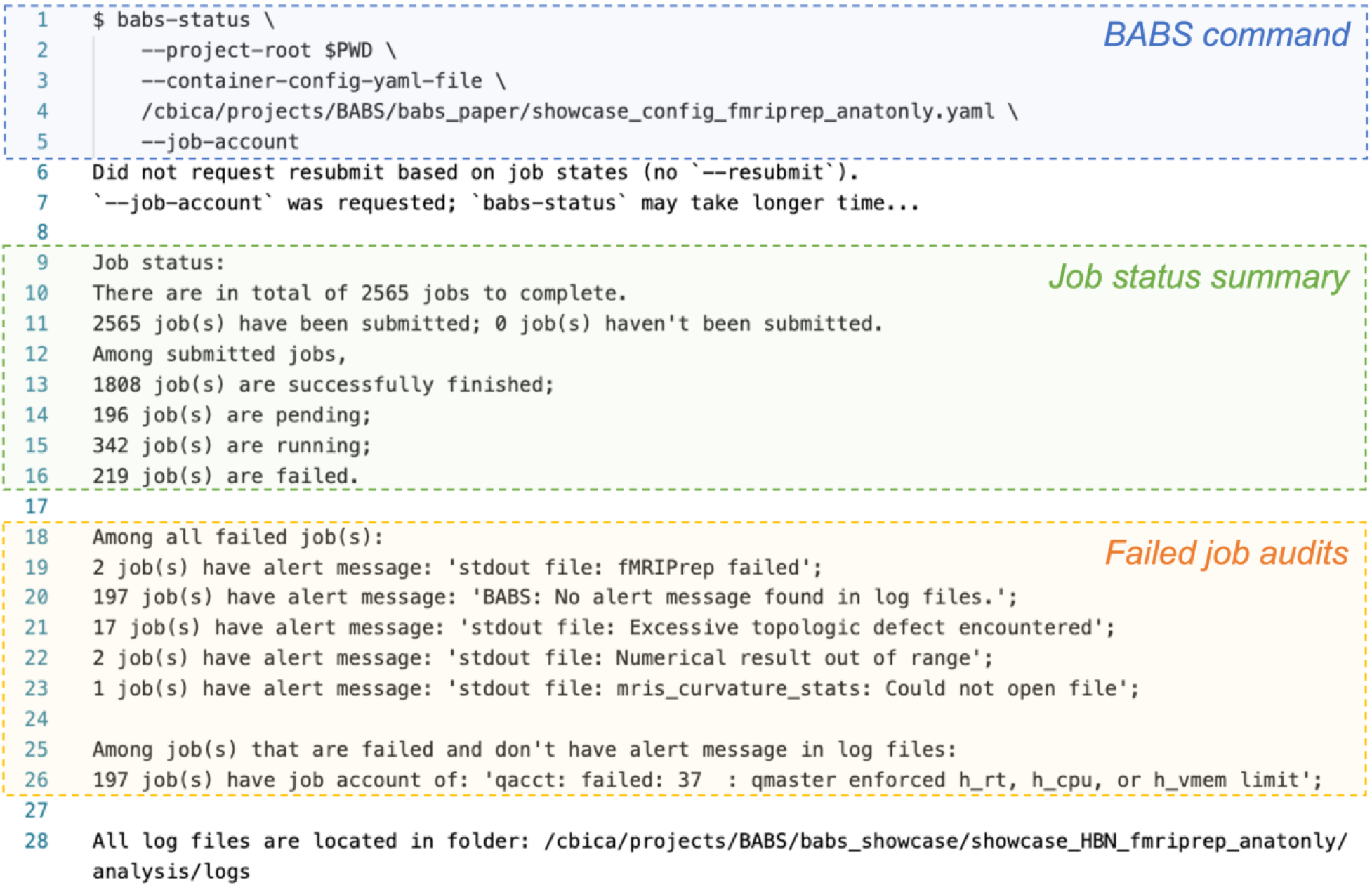
Example `babs-status` command (in blue) and corresponding printed messages during the application of BABS to the HBN dataset. The `babs-status` command was run in the root directory of the BABS project.

Next, to further understand why these jobs failed, and to audit how many jobs had failed due to each reason, `babs-status` performed failed job audits. It first searched for alert messages in log files of these failed jobs that were predefined in the container configuration YAML file (line #46-52, **Figure 4**). These alert messages, such as “`Excessive topologic defect encountered`”, were messages that may be found in log files of failed jobs and may be informative for why a job had failed. `babs-status` identified two failed jobs that had alert message of “`fMRIPrep failed`” in standard output (`stdout`) log file, 17 failed jobs with “`Excessive topologic defect encountered`”, two failed jobs with “`Numerical result out of range`”, and one failed job with “`mris_curvature_stats: Could not open file`” (line #19 and #21-23, **Figure 5**).

Finally, there were 197 failed jobs without predefined alert messages in their log files (line #20, **Figure 5**). This indicates these jobs failed for reasons other than the ones described above. With `--job-account` mode on, `babs-status` further called cluster’s job accounting command to diagnose the reason. All of these 197 failed jobs had failed code `37`: “`qmaster enforced h_rt, h_cpu, or h_vmem limit`” (line #25-26, **Figure 5**). This tells us that these jobs were killed by the cluster because they exceeded resource limits. As we have set job run time limit (`h_rt`) of two days in the container configuration YAML file, these jobs likely failed because they did not finish within two days, indicating they got stuck at one step of the processing pipeline.

As failed job auditing was turned on (i.e., `--container-config-yaml-file` and `--job-account`), this example `babs-status` command at this point finished after around half an hour. Notably, without failed job auditing (using only the job status summary), `babs-status` will only require one to two minutes even for a very large dataset like HBN. Note that the run time of a BABS command also depends on how busy the cluster is, so it is normal for the run time to vary.

After the first round of submissions of all subjects’ jobs, 2,231 jobs successfully finished, and 334 jobs failed. In a large dataset like HBN with heterogeneous data quality, it is expected that jobs failed due to issues such as failed surface reconstruction by FreeSurfer. Sometimes rerunning the jobs may solve the problem. Therefore, we resubmitted these 334 failed jobs using `babs-status` with the argument ‘`--resubmit failed`’. In the end, a total of 2,258 subjects’ jobs successfully finished, and 307 subjects’ jobs failed.

At this time, all successful jobs’ results were on different branches. We applied `babs-merge` to merge them into the mainline branch, resulting in a complete, provenance-tracked DataLad dataset. The results were ready to consume. To check the results, we used DataLad’s command ‘`datalad clone`’ to clone the output RIA to a human readable folder outside the BABS project. After getting into this new folder, we could see 2,258 zip files, each containing results from one subject. By using ‘`datalad get`’, we copied the content of one zip file into the cloned folder. After unzipping this zip file, we successfully got the expected results from the corresponding subject.

To check the data provenance tracked by DataLad in this clone of the results, we used command ‘`git log --oneline`’ to list the Git commit history of the entire workflow of BABS (**Figure 6A**). We further entered the command ‘`git show 09ce675`’ to check the provenance record of the last job shown in the history: commit `09ce675`, for subject `sub-NDARYT155NHX`. This command printed out the machine-readable, re-executable provenance record of this subject’s data processing job, including the exact command used, input BIDS data and container image, results zip file, etc (**Figure 6B**). With this provenance record and the code saved in the output RIA store, data processing of any job can be repeated, making data processing fully reproducible. Here we skip the demonstration of such recomputation, as this has been demonstrated in the FAIRly big framework paper (Wagner et al., 2022).

**Figure 6.**
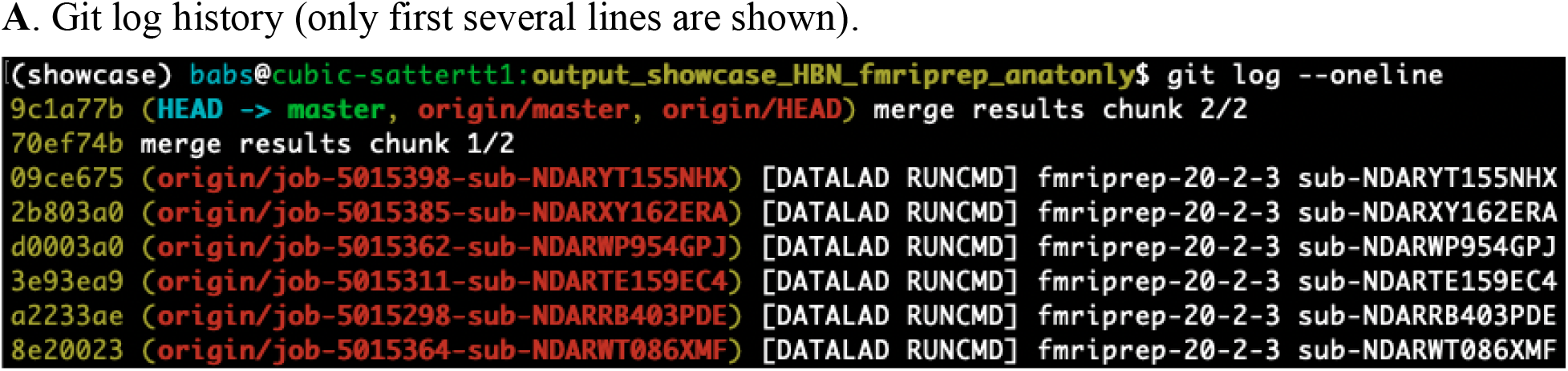
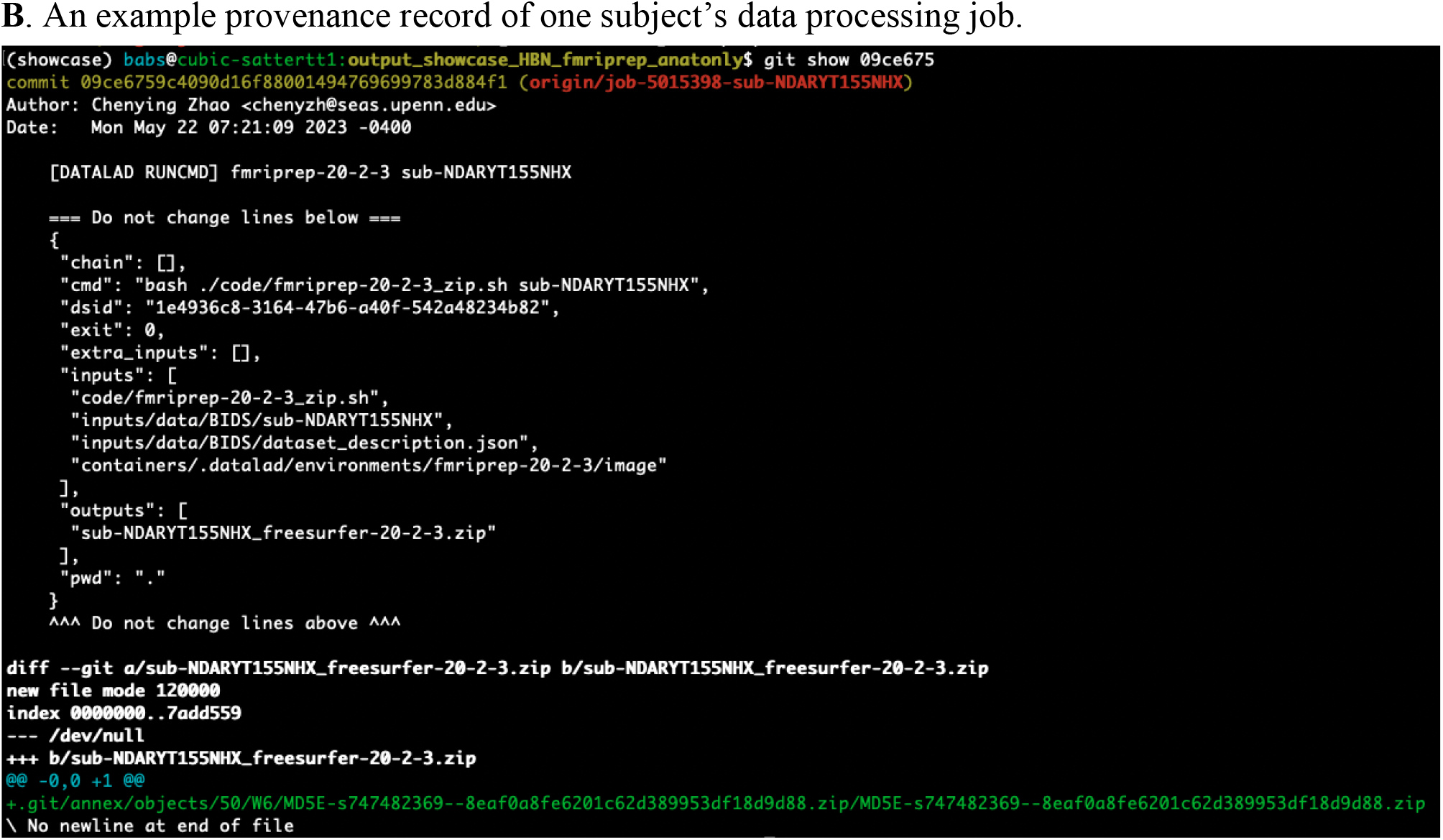
Git log history and example data provenance of the application to the HBN dataset. (**A**) The Git log history of the entire workflow of BABS (only first several lines, i.e., last several commits are shown). (**B**) An example machine-readable, re-executable provenance record of a subject’s data processing job.

## 4. DISCUSSION

BABS was developed to respond to the reproducibility crisis in neuroimaging research and maximize reproducibility in image processing. The full audit trail recorded by DataLad and FAIRly big framework provides complete reproducibility for processing large-scale datasets. As discussed below, BABS allows for reproducible and scalable applications of different BIDS Apps on HPC clusters that are accessible to general users.

### 4.1. Reproducible data analyses

There have been emerging tools for enhancing reproducibility, e.g., Git for code version control, DataLad for data version control, as well as containerized BIDS Apps that are portable and encapsulate all dependent software. However, neuroimaging data analyses often involve various components, including input datasets, software and code, commands and parameters, results, etc. A full audit trail of data processing is essential for complete reproducibility; however, it is often painful to record a full audit trail for complex, heterogeneous, and large datasets. Leveraging DataLad and FAIRly big framework, BABS supports automatic data provenance tracking. This allows users to find detailed data provenance records for results, even from a large-scale dataset. Such detailed records link the results to all other components in neuroimaging data analyses by answering how the results were generated: which input datasets were used, what the exact code, commands, and version of the BIDS App were applied, etc. These detailed records can be retrospectively retrieved when reporting detailed methods or protocol of the study.

It should be noted that, although some platforms such as the Canadian Open Neuroscience Platform (CONP; https://conp.ca/; Harding et al., 2023; Poline et al., 2023) may also use DataLad and containerized software (e.g., BIDS Apps) for enhancing reproducibility, with BABS, it is not necessary for users to upload or share the data on another platform; instead, they can directly process the data on the HPC clusters they have access to.

### 4.2. Scalability to large-scale datasets

Compared to small datasets, large neuroimaging datasets produce more reproducible and replicable findings, especially in the context of the small effect size of many brain-behavior associations (e.g., Marek et al., 2022; Liu et al., 2023). Large sample sizes enhance statistical power; however, their size, complexity and heterogeneity also present challenges in data processing. The challenges include not only data provenance tracking (as discussed above), but also HPC job management. BABS is designed to scale for applications to large-scale datasets in an HPC environment. As shown in the application to the HBN dataset, `babs-submit` and `babs-status` provide simple yet powerful interfaces for users to manage numerous jobs, including job submission, status checking, and auditing of failed job. These features facilitate its applications to large-scale datasets.

### 4.3. Generalizability to different use cases

There are numerous use cases of neuroimaging data processing, including different input BIDS datasets, BIDS Apps, and HPC cluster platforms. It is not efficient for researchers to implement one script per use case; the profusion of resulting code also has the potential to impact reproducibility. Instead, BABS creates (i.e., bootstraps) scripts tailored for different use cases. In this way, maintenance and enhancements are only needed for the source code of BABS, instead of numerous scripts for different use cases. Currently, BABS supports several common use cases, including input BIDS datasets in different forms (one or more; single-session and multiple-session; raw BIDS data and BIDS derivatives), several BIDS Apps (fMRIPrep, QSIPrep, and XCP-D), and SGE and Slurm HPC clusters. Applications to other BIDS Apps are straightforward and can easily be achieved by creating new container configuration YAML files for those BIDS Apps. BABS is also extensible to additional HPC job scheduling systems beyond Slurm and SGE; we have plans to support LSF in the future.

### 4.4. User-friendly and accessible to general users

BABS provides a parsimonious set of programs for reproducible image processing. With BABS, users can apply the FAIRly big framework and reproducibly process large datasets without requiring deep knowledge of DataLad, which many users find challenging. Sanity checks and test jobs included in `babs-check-setup` help users detect problems before real jobs of BIDS Apps start running and fail. Handling numerous jobs on clusters is facilitated by `babs-submit` and `babs-status`; these provide users with concise yet informative messages about the jobs statuses and failed jobs audits. As such, BABS makes the reproducible and large-scale image processing user-friendly and accessible to both beginning and advanced users.

### 4.5. Limitations and future directions

It should be noted that BABS has several limitations. First, BABS capitalizes on and supports the standard BIDS format and BIDS Apps, and is not currently compatible with data in other formats, or neuroimaging software or containers that are not compliant with the requirements of BIDS Apps. However, BIDS data and BIDS Apps are in wide current use. Second, currently BABS is designed to be applied on HPC systems, and is not compatible with cloud-based computing platforms (e.g., Amazon Web Services [AWS]), local computers where a job scheduling system or Singularity software is not installed, or computing nodes without job scheduling systems. Compatibility to these systems may be considered in the future. Third, BABS currently only has command-line interfaces and does not have a graphical user interface (GUI). However, users who use HPC clusters should have basic skills of using command-line programs in terminals. In addition, given that HPC clusters often have limited bandwidth for graphical data transmission, GUI may not be an optimal choice for BABS.

BABS’s continued development is an ongoing, collaborative effort. In the future, several enhancements on the roadmap may be added to BABS. For example, currently the data from the input BIDS dataset(s) should be fixed for a certain BABS project. This may not be suitable for studies where data is still being collected. In the future, we may add functionality to BABS so that it can accept new subjects and/or sessions easily. In addition, as listed in the NMIND checklist (see **Appendix**; Kiar et al., 2023), although BABS has achieved Bronze tiers for documentation, infrastructure, and testing according NMIND standards, there are still several features that remain un-implemented. We have listed these un-implemented features and potential enhancements on the roadmap in a milestone on GitHub: https://github.com/PennLINC/babs/milestone/3. We welcome contributions from researchers in the broad community via pull requests at BABS GitHub repository (https://github.com/PennLINC/babs); developer documentation can be found at: https://pennlinc-babs.readthedocs.io/en/stable/developer.html.

### 4.6. Conclusion

BABS is a Python package that provides a reproducible and scalable workflow for large-scale BIDS data analysis using BIDS Apps. It provides a parsimonious set of user-friendly programs to allow for the generalizable implementation of the FAIRly big workflow on HPC systems. Taken together, BABS facilitates reproducible neuroimaging research at scale.

## AUTHOR CONTRIBUTIONS (CRediT statement)

**Chenying Zhao:** Conceptualization, Formal analysis, Investigation, Methodology, Software, Validation, Visualization, Writing – original draft, Writing – review & editing. **Dorota Jarecka:** Formal analysis, Funding acquisition, Investigation, Methodology, Software, Validation, Writing – review & editing. **Sydney Covitz:** Conceptualization, Data curation, Formal analysis, Methodology, Software, Validation, Writing – review & editing. **Yibei Chen:** Validation, Writing – review & editing. **Simon B. Eickhoff:** Funding acquisition, Writing – review & editing. **Damien A. Fair:** Resources, Writing – review & editing. **Alexandre R. Franco:** Investigation, Resources, Writing – review & editing. **Yaroslav O. Halchenko:** Funding acquisition, Methodology, Software, Writing – review & editing. **Timothy J. Hendrickson:** Resources, Writing – review & editing. **Felix Hoffstaedter:** Methodology, Software, Writing – review & editing. **Audrey Houghton:** Resources, Writing – review & editing. **Gregory Kiar:** Funding acquisition, Investigation, Resources, Writing – review & editing. **Austin Macdonald:** Software, Writing – review & editing. **Kahini Mehta:** Writing – review & editing. **Michael P. Milham:** Funding acquisition, Investigation, Resources, Writing – review & editing. **Taylor Salo:** Methodology, Software, Writing – review & editing. **Michael Hanke:** Funding acquisition, Methodology, Software, Writing – review & editing. **Satrajit S. Ghosh:** Funding acquisition, Methodology, Project administration, Software, Supervision, Writing – review & editing. **Matthew Cieslak:** Conceptualization, Data curation, Methodology, Project administration, Software, Supervision, Visualization, Writing – review & editing. **Theodore D. Satterthwaite:** Conceptualization, Funding acquisition, Methodology, Project administration, Software, Supervision, Visualization, Writing – original draft, Writing – review & editing.

## ACKNOWLEDGEMENTS

This study was supported by grants from the National Institutes of Health: R01MH112847 (T.D.S.), R01MH120482 (T.D.S.), R37MH125829 (T.D.S.), R01EB022573 (T.D.S.), R01MH113550 (T.D.S.), RF1MH116920 (T.D.S.), P41EB019936 (S.S.G., D.J., Y.O.H.), RF1MH130859 (G.K., M.P.M.). S.B.E. and M.H. acknowledge funding by the European Union’s Horizon 2020 Research and Innovation Program (grant agreement 945539 (HBP SGA3)), and the Deutsche Forschungsgemeinschaft (DFG, SFB 1451 (431549029) & IRTG 2150 (269953372)). M.H. and Y.O.H. were supported by co-funding from the US National Science Foundation (NSF 1912266) and German Federal Ministry of Education and Research (BMBF 01GQ1905). Additional support was provided by the AE Foundation and the Penn/CHOP Lifespan Brain Institute.

## APPENDIX

### NMIND checklist

The NMIND checklist is a tool to evaluate the scientific software against coding standards proposed by NMIND consortium (Kiar et al., 2023). The full checklist can be found here: https://www.nmind.org/standards-checklist/. Here we show our Python package BABS’s performance on this checklist (checklist version: 1.0.0; evaluated on August 2nd, 2023). This checklist includes three domains: documentation, infrastructure, and testing. The assessment for each domain could land on one of these three tiers: Bronze, Silver, Gold. The achieved items are labeled with checkmarks “√”, while not achieved ones are labeled with open circles “○”.

It should be noted that, current checklist is based on the latest version of BABS when the paper is submitted. As BABS is under continued development, after more features are added, more checks may be achieved than those listed below, and BABS may even reach a higher tier.

### Domain #1: Documentation

<u>Bronze tier (9 out of 9 items have been achieved):</u>

√ Landing page (e.g., GitHub README, website) provides a link to documentation and brief description of what program does
√ Documentation is up to date with version of software
√ Typical intended usage is described
√ An example of its usage is shown
√ Document functions intended to be used by users (i.e., public function docstring / help coverage ≥ 10%)
√ Description of required input parameters for user-facing functions with reasonable description of inputs (i.e., “NIfTI of brain mask in MNI” vs. “An image file”)
√ Description of output(s)
√ User installation instructions available
√ Dependencies listed (i.e., external and within-language requirements)

<u>Silver tier (4 out of 7 items have been achieved):</u>

√ All items from bronze tier
○ Background/significance of program
√ One or more tutorial to showcase the multiple of the program’s usages (i.e., if program has multiple usages)
○ Any alternative usage that is advertised is thoroughly documented
√ Thorough description of required and optional input parameters
√ Document public functions (i.e., public function docstring / help coverage ≥ 20%)
○ A statement of supported operating systems / environments (i.e., could be a container recipe)

<u>Gold tier (3 out of 8 items have been achieved):</u>

√ All items from bronze tier
○ All items from silver tier
○ Continuous integration badges in README for build status
√ Continuous integration badges in README for tests passing
○ Continuous integration badges in README for coverage
√ Document functions, classes, modules, etc. (i.e., public + private docstring / help coverage ≥ 40%)
○ Has a documented style guide
○ Maintenance status is documented (e.g., expected turnaround time on pull requests, whether project is maintained)

### Domain #2: Infrastructure

<u>Bronze tier (7 out of 7 items have been achieved):</u>

√ Code is open source
√ Package is under version control
√ Readme is present
√ License is present
√ Issues tracking is enabled (i.e., either through GitHub or external site)
√ Digital Object Identifier (DOI) points to latest version (e.g., Zenodo)
√ All documented installation instructions can be successfully followed

<u>Silver tier (2 out of 4 items have been achieved):</u>

√ All items from bronze tier
○ Issue template(s) available (i.e., information requested by developers)
√ Continuous integration runs tests
○ No excessive files included (i.e., unused files / cache; e.g., .gitignore)

<u>Gold tier (3 out of 7 items have been achieved):</u>

√ All items from bronze tier
○ All items from silver tier
√ Continuous integration builds packages
○ Continuous integration validates style
○ Journal of Open Source Software submission
√ Contribution guide present
○ Code of Conduct present

### Domain #3: Testing

<u>Bronze tier (2 out of 2 items have been achieved):</u>

√ Provide / generate / point to test data
√ √ Provide instructions for users to run tests that include instructions for evaluation for correct behavior

<u>Silver tier (2 out of 3 items achieved):</u>

√ All items from bronze tier
√ Some form of testing suite present
○ Test coverage > 50%

<u>Gold tier (2 out of 4 items have been achieved):</u>

√ All items from bronze tier
○ All items from silver tier
○ Test coverage > 90%
√ Benchmarking information is provided for examples

